# Recalibrating the cerebellum DNA methylation clock: implications for ageing rates comparison

**DOI:** 10.1101/2022.05.16.492116

**Authors:** Yucheng Wang, Olivia A. Grant, Xiaojun Zhai, Klaus D. McDonald-Maier, Leonard C. Schalkwyk

**Author notes:** **Correspondence**: Xiaojun Zhai and Leonard C. Schalkwyk.

## Abstract

**Background:** DNA methylation (DNAm) based age clocks have been studied extensively as a biomarker of human ageing and risk factor for age-related diseases. Despite different tissues having vastly different rates of proliferation, it is still largely unknown whether they age at different rates. It was previously reported that the cerebellum ages slowly, however, this claim was drawn from a single clock using a small sample size and so warrants further investigation.

**Results:** We collected the largest cerebellum DNAm dataset (N=752). We found the respective epigenetic ages are all severely underestimated by six representative DNAm age clocks, with the underestimation effects more pronounced in the four clocks whose training datasets do not include brain-related tissues. We identified 613 age-associated CpGs in the cerebellum, which accounts for only 14.5% of the number found in the middle temporal gyrus from the same population (N=404), of which only 201 CpGs are both age-associated in the two tissue types. We built a highly accurate age prediction model for the cerebellum named CerebellumClock_specific_ (Pearson correlation=0.941, MAD=3.18 years). Furthermore, based on the 201 age-associated CpGs, we built two other clocks CerebellumClock_common_ and CortexClock_common_ for the cerebellum and non-cerebellar brain cortex tissues separately, they both support that the cerebellum has a relative lower DNAm ageing rate.

**Conclusions:** The large underestimation for the cerebellum by previous clocks mainly reflects the improper usage of the age clocks. There exist strong and consistent ageing effects on the cerebellar methylome despite the cerebellum having unique age-dependent methylome changes. The DNAm clock based ageing rates comparisons are valid only upon models constructed on a small group of CpGs, therefore, more evidence is required to support the idea that different DNAm ageing rates represent different biological ageing rates.

## 1 BACKGROUND

Ageing is characterized by progressive loss of cellular functions, leading to increased risk of morbidity and mortality [1]. A significant challenge in the ageing field is how to accurately measure ageing. Further investigation of ageing biomarkers will not only increase our knowledge of the mechanisms of ageing, but also facilitate monitoring the various interventions for improving human healthspan and rejuvenation experiments. In the past decades, a variety of ageing biomarkers, such as telomere attrition [2], DNA methylation (DNAm) changes [3, 4] and alterations in gene expression [5, 6] and metabolite concentration [7, 8], have attracted even more attention and were used to build age estimators or age clocks to measure the biological age [9]. Among them, age clocks based on DNAm changes, also called epigenetic clocks, were demonstrated to be the most accurate and robust age estimators, they are the most promising ageing biomarkers that can be applied to individuals [10]. Age-related DNAm changes are widespread across the genome, throughout the life course [11, 3, 12, 13] and exist in a wide variety of tissues [4, 14].

Since 2013, many DNAm-based age clocks have been published. Among them, Hannumn’s clock [3] and Horvath’s clock [4] are two first and most widely used DNAm clocks, especially Steve Horvath first demonstrated that a relatively accurate age prediction (median absolute error 3.6 years) is possible for multiple distinct tissues via a single linear model that includes only a small number of CpGs [4]. Different tissues may have distinct DNA methylation profiles, therefore many tissue-specific clocks have been developed and demonstrated better age prediction performance than the multi-tissue clock, tissue-specific clocks have been developed for buccal cells [15], brain cortex [16], skin [17] and so on. Age acceleration, a popular concept that defined as the difference between predicted DNAm age and the chronological age, derived from the Hannnum’s clock or Horvath’s clock has been documented to be associated with a variety of age-related conditions and diseases [18, 19, 20, 21]. However, Zhang et al. reported that the association between age acceleration and mortality decreased to non-significant with increased accuracy of chronological age prediction of age models [22]. Meanwhile, instead of directly regressing on the chronological age, two other DNAm clocks—PhenoAge [23] and GrimAge [24] which were regressed on estimated phenotypic age and mortality risk respectively, reported to better predict lifespan and healthspan than previous chronological age clocks.

Ageing is generally considered a gradual process that happens to the body as a whole. It is still an open question whether different organs/tissues have different ageing rates. Furthermore, how can we truthfully compare the ageing rates between different tissues? Horvath’s pan-tissue clock gives excellent accuracy in estimated DNAm age for many different cells and tissues [4], which may suggest that those different cell and tissue types may have similar ageing rates. In 2015, Horvath et al. claimed that the cerebellum ages slower than many other parts of the human body based on the observations that the DNAm age of the cerebellum is much lower than other tissues based on the pan-tissue clock [25]. In addition, Horvath and his colleagues also claimed that women’s breast tissues have a relative higher DNAm ageing rate [4, 26]. If it is true that some tissues have significant different DNAm ageing rates than other tissues, then we can go further to identify what drives the difference. This is a very important angle to understand the behind mechanisms of the age-associated DNA methylation changes. Even though there have been reported many strong age-associated CpG sites, there is still very little known about the underlying mechanisms that drive age-associated DNA methylation changes [27, 28, 29].

In recent years, many more cerebellum DNA methylation samples have become publicly available and many diverse DNAm age clocks have also been developed [30]. We seek to examine the claim that the cerebellum ages slowly with a much larger dataset and try to find out the mechanisms. To achieve that, we first collect the largest cerebellum DNA methylation sample dataset, then compare their estimated epigenetic ages from six representative DNAm age clocks; secondly, we perform age EWAS for two brain tissues (i.e. cerebellum and middle temporal gyrus) separately on a large elderly population (n = 404), to reveal the distinct age-associated DNAm changes of the cerebellum. Lastly, we construct cerebellum specific clocks and further examine the claim that the cerebellum ages slower.

## 2 RESULTS

### 2.1 Characteristics of the DNAm cerebellum datasets

The cerebellum is a structure of the hindbrain, which plays a vital role in motor control [31]. Unlike peripheral tissues, such like blood or saliva, that can be non-invasively and repeatedly sampled, cerebellum samples are often collected from the post-mortem subjects, as a result, there is a very limited number of DNAm cerebellum samples available. After rigorous searching on the Gene Expression Omnibus (GEO) database, where publicly available DNAm datasets are often deposited, we found a total of 6 datasets, that each includes more than ten cerebellum samples measured by Illumina 450k or EPIC array. After rigorous quality control (see methods), 752 cerebellum samples remained and were used for downstream analysis. The biggest contributor for the final big cerebellum dataset is from GSE134379 [32], which contains 404 cerebellum samples. As cerebellum samples were from postmortem subjects, 90% of the collected samples were from individuals aged above 60 years old, with the median age at 80 years old. More detailed age, sex and disease distribution information for each dataset are listed in Table 1. The DNAm microarray data from those datasets were originally produced to investigate disease-associated methylomic variations in the brain regions, especially for Alzheimer’s disease and Schizophrenia. As a result of this, our collected cerebellum samples include 333 samples with normal health status, 398 samples with Alzheimer’s disease and 21 with Schizophrenia. The cerebellum is a relatively protected region, unlike other brain regions (such as prefrontal cortex), there generally are no significant AD-associated differences in the cerebellum [33, 34, 35]. Therefore, we included all these cerebellum samples, even those with disease diagnosis, for downstream analysis and also for the following cerebellum DNAm age clock construction.

**TABLE 1.** Characteristics the clean cerebellum samples from six datasets

| ID | Number | Female, Male | Age: mean (range) | Disease group | Reference |
| --- | --- | --- | --- | --- | --- |
| GSE134379 | 404 | 200, 204 | 83.7 (54-103) | Alzheimer: 225<br>Normal: 179 | [32] |
| GSE59685 | 111 | 64, 47 | 83.9 (40-105) | Alzheimer: 59<br>Normal: 52 | [34] |
| GSE105109 | 95 | 41,54 | 81.2 (58-99) | Alzheimer: 67<br>Normal: 28 | [36] |
| GSE125895 | 66 | 32, 34 | 67.3 (51.8-92.3) | Alzheimer: 24<br>Normal: 42 | [35] |
| GSE61431 | 44 | 16, 28 | 61.6 (25-96) | Schizophrenia: 21<br>Normal: 23 | [37] |
| GSE72778 | 32 | 21, 11 | 83.2 (15-114) | Alzheimer: 23<br>Normal: 9 | [38] |

### 2.2 Severe underestimation for cerebellum samples by various DNAm age clocks

Since 2013, many specialized and robust DNAm based clocks have been reported. As recently suggested by Liu et al., those different clocks may have captured different biological processes of ageing considering their overall weak associations in the estimated DNAm age deviations [39]. Inspired by this, we investigated the DNAm ages of our collected 752 cerebellum samples predicted by six representative clocks: Hannum’s whole blood clock (Hannum2013) [3] and Horvath’s pan-tissue clock (Horvath2013)[4] are the two most widely used DNAm age clocks and especially Horvath2013 is reported to work well across many different tissue and cell types; Horvath’s blood&skin clock (Horvath2018) [27] is another multi-tissue clock and was reported to outperform Horvath2013 across several tissues; Levine’s PhenoAge clock (Levine2018) [23] was not directly regressing on chronological age and reported better prediction performance for all-cause mortality than other chronological age regressed clocks; Zhang’s blood clock (Zhang2019) [22] is reported the most accurate and robust age prediction model for blood samples; Shireby’s brain cortex clock (Shireby2020) [16] is a brain cortex specific clock and provides far better age predictions than other clocks in brain cortex tissues.

As shown in Figure 1, almost all of the cerebellum samples are severely underestimated—they are all distributed below the diagonal lines. Hannum2013, Levine2018 and Zhang2019 are three age clocks trained almost exclusively on blood samples, the root-mean-square deviations (RMSDs) of their predictions are all very large (above 40 years), with Pearson correlations (r) ranging from 0.182 in Levine2018 and 0.56 in Zhang2019; Horvath2018 is a multi-tissue clock that was trained on eight different tissues/cells not including brain-related tissues, it gave a similar prediction trend for cerebellum samples as the three blood clocks—large deviations (RMSD=66.9 years) and low correlation (*r*=0.452). The underestimation effect is less apparent for Horvath2013 and Shireby2020, their RMSDs are just above 20 years and Pearson correlation 0.699 in Shireby2020 and 0.694 in Horvath2013. This may be because Horvath2013 is a pantissue clock which included several different brain-related tissues including 282 cerebellum samples in its training dataset [4] while Shireby2020 was trained exclusively on brain cortex. It should be noted, the regression line of the predicted DNAm age against the chronological age in Horvath2013 and Shireby2020 both suggest that the cerebellum samples from young individuals aged below 30 years old are very likely to be overestimated. Overall, the systematic severe underestimation trend and the smaller than 1 value of the regression coefficient in all six clocks appear to support the claim that the cerebellum ages slowly.

**FIGURE 1.**
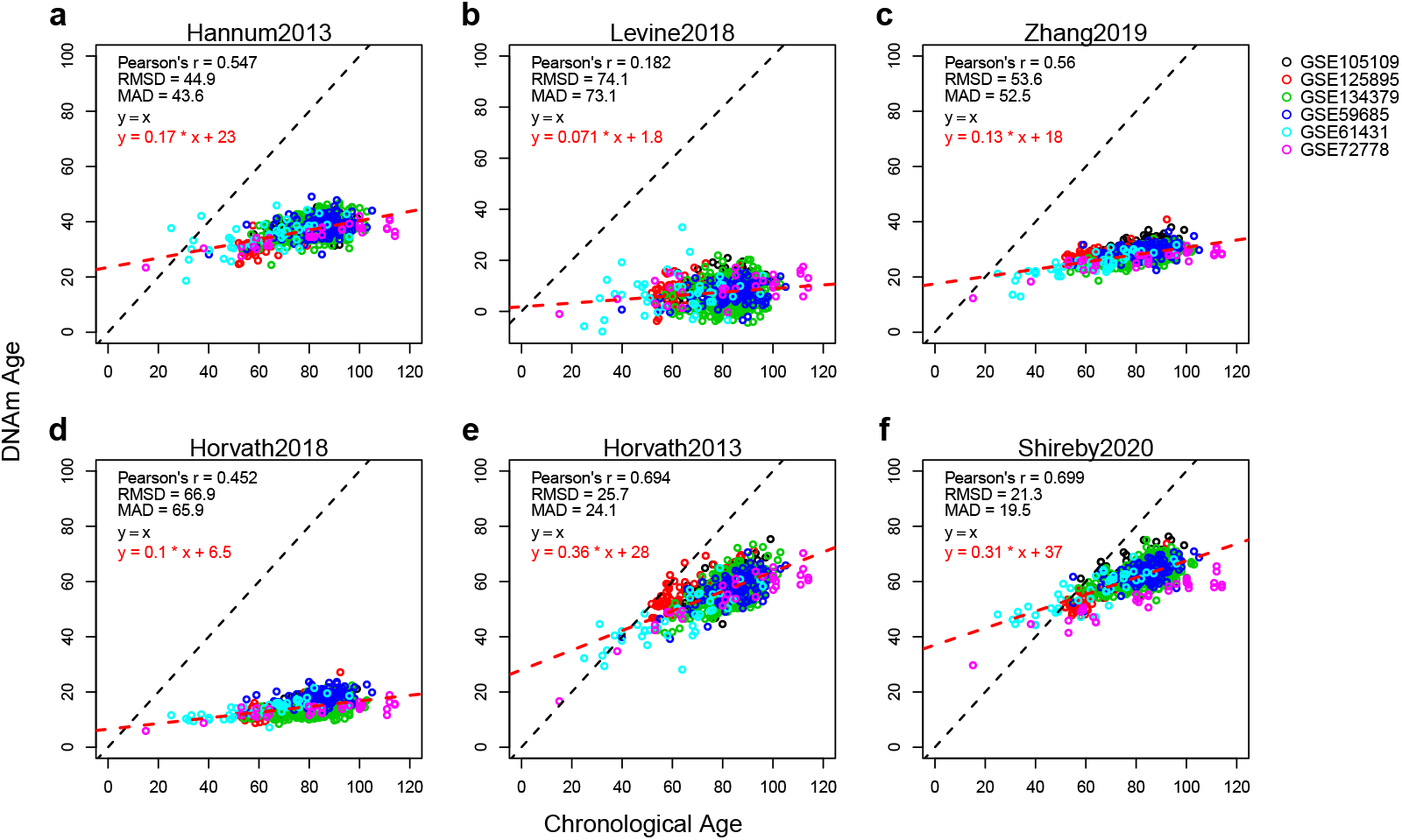
The cerebellum samples are severely underestimated by the six representative DNAm clocks. Each subplot illustrates results from different clocks: (**a**) Hannum2013, (**b**) Levine2018, (**c**) Zhang2019, (**d**) Horvath2018, (**e**) Horvath2013 and (**f**) Shireby2020. The colorful dots represent 752 cerebellum samples from six independent datasets, with different colors representing different datasets. The x-axis is chronological age and the y-axis is the estimated DNAm age. The black dashed line represents the identical diagonal line between chronological age and DNAm age, the red dashed line represents the regression line derived from regressing the DNAm age against the chronological age. RMSD: root mean squared deviation; MAD: mean absolute deviation.

### 2.3 Distinct age-associated methylomic changes of the cerebellum

We went further to investigate the underlying reasons why the cerebellum is systematically underestimated by the six age clocks. We hypothesized that, if the cerebellum truly ages slower than most other brain tissues, then due to a smaller ageing effect, there would be a much smaller number of CpGs passing the same cutoff to be identified as age-associated and even those captured age-associated CpGs would mostly exhibit a smaller rate of age-associated methylated changes. Inspired by this, we carried out two epigenome-wide association studies (EWAS) on age for the cerebellum (CBL) and the middle temporal gyrus (MTG) separately, based on the same dataset GSE134379 [32] which includes DNA methylation microarray samples of the two brain regions for every subject from a large elderly population (n=404).

We obtained a total of 4,213 significant (adjusted P-value ≤0.01) age-associated CpGs in MTG, in contrast, only 613 CpGs were found to be age-associated in CBL (Figure 2a, 2b and Supplementary 1). This once again seemingly supports the claim that the cerebellum has a slower ageing rate. However, comparing the top 613 significant age-associated CpGs between the two tissues, only 86 (14%) CpGs are shared by both tissues (Figure 2c). Even looking at all the age-associated CpGs in both tissues, only 32.8% (201) of that CpGs in CBL were also been captured in MTG. More interestingly, when looking at the direction of effect, CBL and MTG showed very different patterns in their age-associated CpGs. The CBL-only group have similar numbers of positive and negative age association. In contrast, more than three quarters (76%) of the MTG-only CpGs gain methylation with ageing. Surprisingly, the majority (94%) of the age-associated CpGs shared between tissues are positively associated (Figure 2d). It should be also noted, among the shared 201 CpGs, there is a slightly higher proportion (56%) of CpGs which has a larger effect size in age association in CBL than that in MTG (Figure 2d). Collectively, even though the cerebellum has a much smaller number of age-associated CpGs, the smaller number seems not directly due to a smaller ageing effect, as the age-associated CpGs in the cerebellum have only a small intersection with that in MTG, but may contribute to the unique age response of the cerebellum.

**FIGURE 2.**
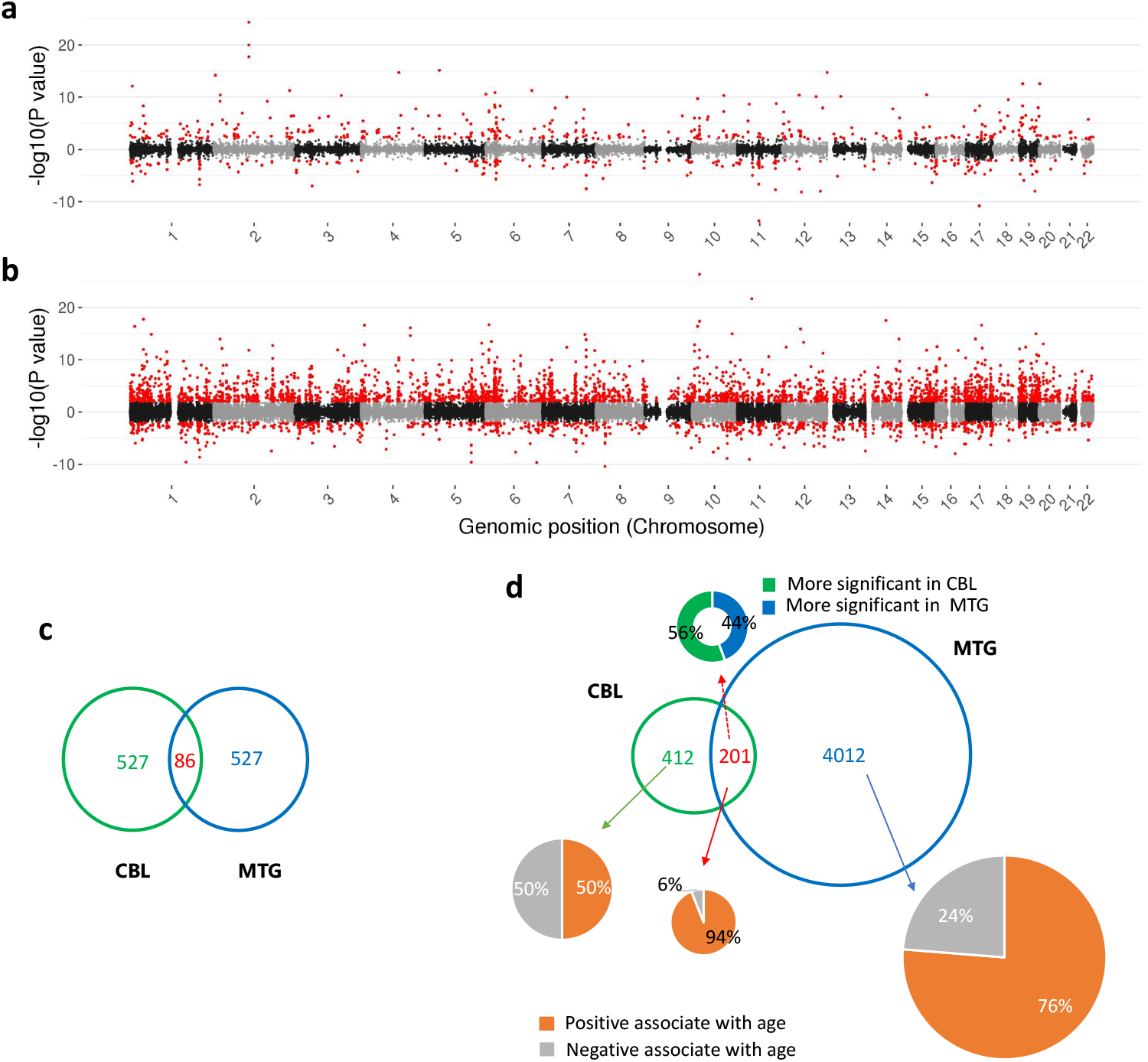
Comparison of age-associated methylation change between the cerebellum (CBL) and the middle temporal gyrus (MTG). Manhattan plots illustrate the age EWASs results of (**a**) CBL and (**b**) MTG. Red dots denote significant age-associated CpGs (adjusted P-value ≤ 0.01). (**c**) Venn plot shows the unique and shared number of the top 613 most significant age-associated CpGs in CBL and MTG. (**d**) Comparison of the age-associated CpGs in CBL and MTG. The three pie charts illustrate the proportions of CpGs gain methylation (positive associate with age) or lose methylation (negative associate with age) with age. The upper doughnut chart shows the proportions of CpGs exhibiting higher age association in CBL or MTG.

Gene ontology analyses showed several enriched terms for the MTG-specific CpGs and Cerebellum-specific CpGs (Supplementary 2) which included terms related to chromatin such as DNA binding, nucleosome assembly and negative regulation of transcription by RNA polymerase II. The MTG-specific CpGs were enriched for pathways such as telomere organisation, noradrenergic neuron differentiation and dopaminergic neuron differentiation. Telomere shortening and neuron dedifferentiation are both characteristics of healthy ageing so it is interesting that these pathways are enriched for the MTG specific CpGs but not for the cerebellum specific CpGs. The enriched GO terms for the cerebellum specific CpGs were related to molecular functions such as DNA binding activity.

### 2.4 The DNAm age clock for cerebellum

#### 2.4.1 Training the cerebellum specific DNAm clock

Our analyses in the previous section have clearly demonstrated that the six representative DNAm age clocks, including the pan-tissue clock and the brain cortex clock, all severely underestimated in cerebellum samples. In addition, we have shown that the cerebellum has a much smaller number of age-associated CpGs. Then we went further to find out whether it is possible to build an accurate age prediction model for the cerebellum.

We trained a cerebellum-specific age model, named CerebellumClock_specific_, by regressing the beta values of the 613 age-associated CpGs from the 752 clean cerebellum samples against their corresponding chronological ages via the Elastic Net penalized linear regression algorithm [40]. The prediction performance of the model was measured by leave-one(dataset)-out cross validation (see Methods). As shown in Figure 3a, the cross validation results demonstrate that the trained cerebellum age models yield very good age predictions for nearly all cerebellum datasets, except that most of the elderly subjects in GSE72778 were relatively underestimated. The overall Pearson correlation is above 0.94, with RMSD at 4.26 years and MAD is 3.18 years. The accurate age prediction performance of the cerebellum age model demonstrates that there is persistent and significant ageing process undergoing in the cerebellum tissues.

**FIGURE 3.**
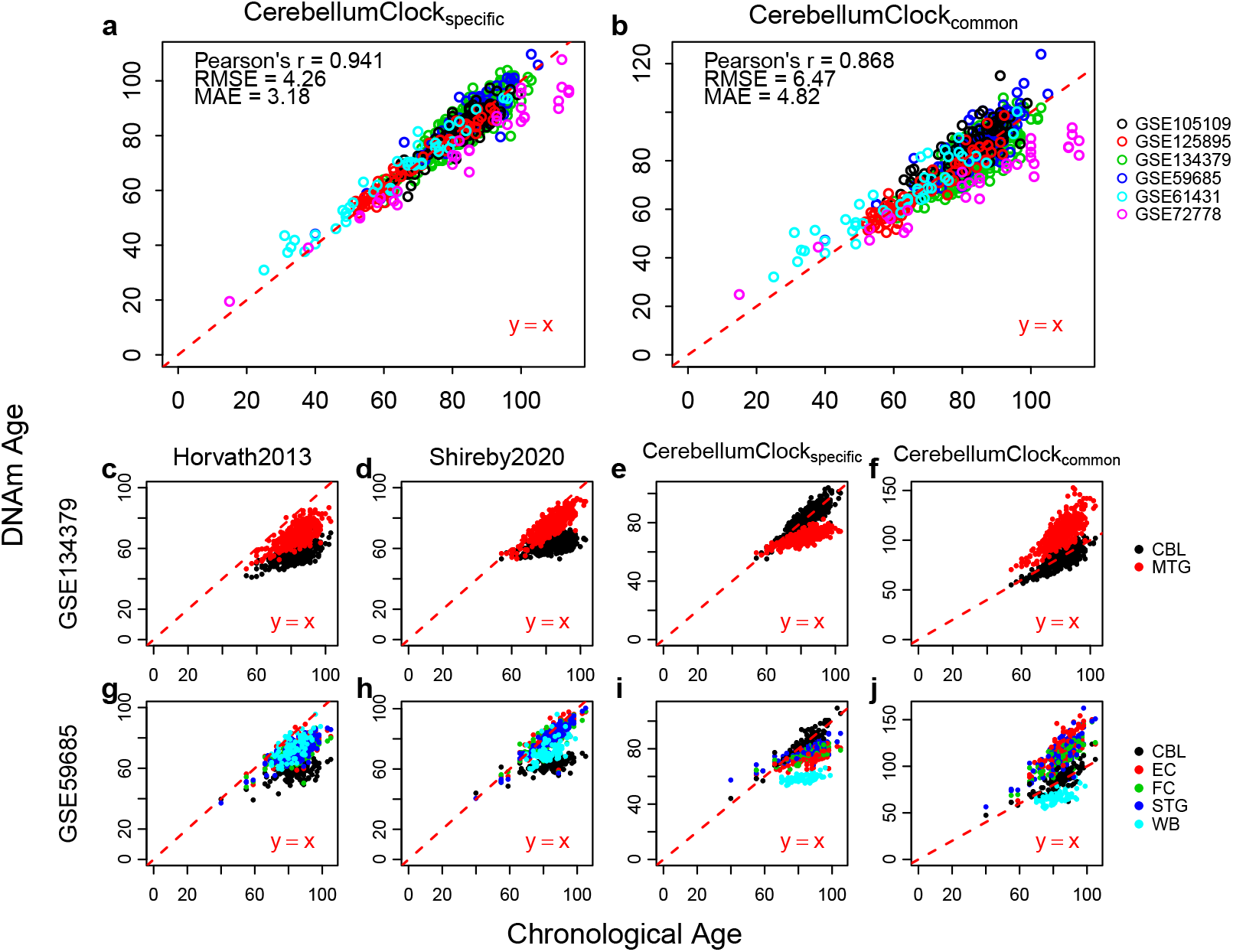
The cerebellum age clocks and their applications in other tissues. The leave-one(dataset)-out cross validation evaluate the age prediction performance of (**a**) CerebellumClock_specific_ and (**b**) CerebellumClock_common_ cerebellum samples. Subplots (**c**), (**d**), (**e**) and (**f**) compare the DNAm age of CBL and MTG estimated by Horvath2013, Shireby2020, CerebellumClock_specific_ and CerebellumClock_common_ respectively. Similarly, Subplots (**g**), (**h**), (**i**) and (**j**) compare the DNAm age of five different tissues estimated by the same four clocks. CBL: cerebellum, MTG: middle temporal gyrus, EC: Entorhinal Cortex, FC: Frontal Cortex, STG: Superior Temporal Gyrus, WB: Whole Blood.

#### 2.4.2 Applying the cerebellum clocks in other tissues

To further test the claim that cerebellum ages slower, we made another hypothesis: other tissues, including brain cortex and blood, would be significantly overestimated for their DNAm ages when measured by the cerebellum clock. To test the hypothesis, we then applied CerebellumClock_specific_ along with Horvath2013 and Shireby2020 in two separate datasets: GSE134379 and GSE59685, which both include cerebellum samples and samples of other tissues from the same subject. As expected, the cerebellum samples were apparently underestimated compared to other tissues by Horvath2013 and Shireby2020 in both GSE134379 and GSE59685 (Figure 3). Interestingly, even though blood was also not included in the training set of Shireby2020, the predicated DNAm ages of blood samples in GSE59685 are still much higher than their counterparts in the cerebellum tissue (Figure 3h). Surprisingly, when measured by our cerebellum clock, the non-cerebellar samples were actually underestimated rather than overestimated compared to the cerebellum tissue (Figure 3i). This finding against our previous expectation, we speculate the underestimation effect for other tissues by the cerebellum clock may rather imply that the age model is working poorly in non-cerebellar tissues. We could observe a trend that the cerebellum clock seems to overestimate these non-cerebellar samples whose ages are below 60 years old (Figure 3e and 3i). This is further confirmed by looking at the overestimation facts for cortex tissues from young subjects by the cerebellum clock (Supplementary Figure 1). The penalized regression algorithm selected 275 age-associated CpGs from the 613 age-associated CpGs in the cerebellum, where the majority of them (73%) are on the CBL-only list, meaning they do not exhibit significant age associations in MTG. Thus the observed apparent underestimation effect for those non-cerebellar samples is not biologically meaningful, instead indicating the improper usage of age models.

### 2.5 Ageing rates comparison

The first model thus captures cerebellum-specific age-related changes. We then trained another cerebellum age model with the same regression algorithm and the same training samples except the input CpG set was restricted to the 201 CpGs that are age-associated in both CBL and MTG. The leave-one(dataset)-out cross validation demonstrated that the new cerebellum clock, named CerebellumClock_common_, still gives very good age predictions for those cerebellum samples (Figure 3b). Notably, the new cerebellum clock CerebellumClock_common_ substantially overestimated the non-cerebellar brain tissues in both GSE134379 and GSE59685, though the blood samples in GSE59685 were still underestimated (Figure 3f and 3j).

To further confirm the overestimation effect for non-cerebellar brain tissues by the new cerebellum clock, we applied the CerebellumClock_common_ in another two independent datasets which combined includes a large number of samples from non-cerebellar brain tissues and has a wide age range (20~100 years old). The results in Figure 4a demonstrate that CerebellumClock_common_ substantially overestimates almost all the non-cerebellar brain tissues. The overall Pearson correlation reached 0.951, indicating the new cerebellum clock has also captured the strong ageing effect on the methylome in those tissues. More importantly, the slope of the regression line obtained from regressing the predicted DNAm age against the chronological age is greater than 1 (1.2), indicating that these non-cerebellar tissues have a higher ageing tick rate than cerebellum.

**FIGURE 4.**
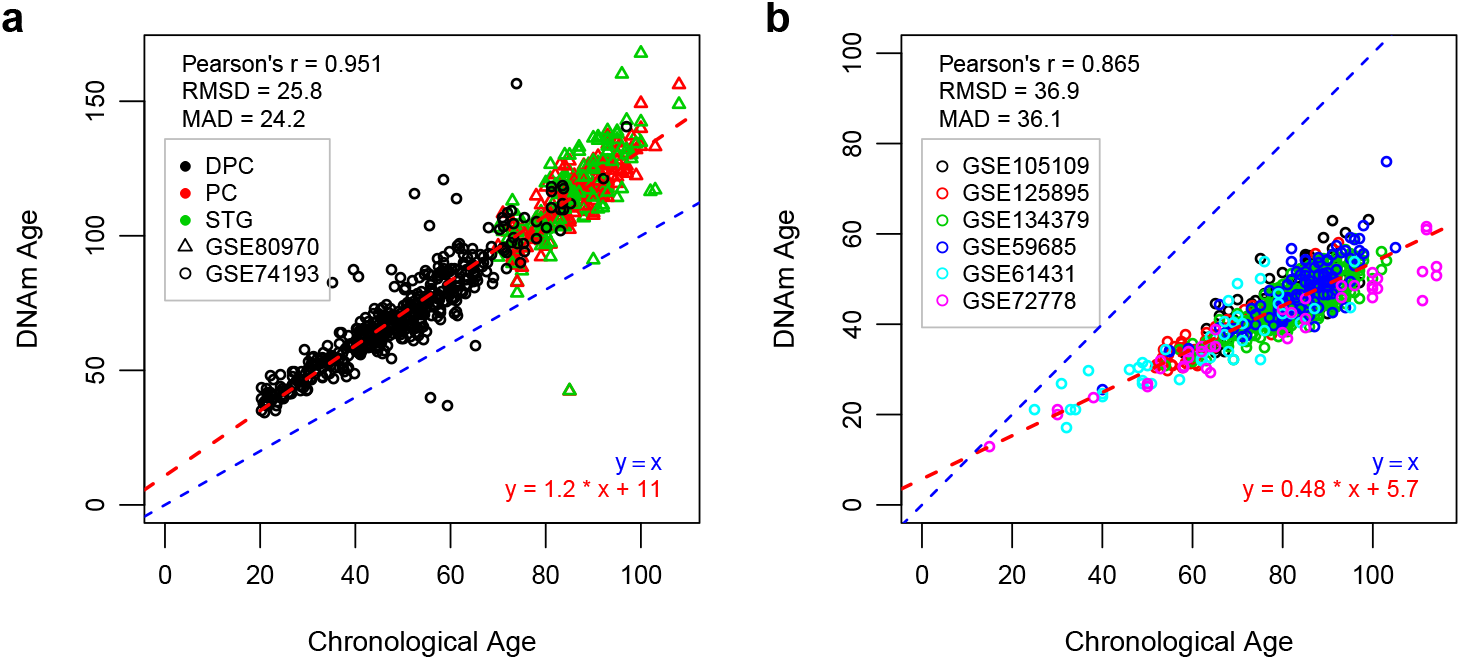
(a) The DNAm ages of three brain tissues (DPC: Dorsolateral Prefrontal Cortex, PC: Prefrontal Cortex, STG: Superior Temporal Gyrus) from two datasets are systematically overestimated by CerebellumClock_common_. (**b**) the cerebellum samples are all hugely underestimated by the CortexClock_common_. The CerebellumClock_common_ and CortexClockcommon were trained from the same set of CpGs (n=201) but in different tissues. The blue dashed line represents the identical diagonal line between chronological age and DNAm age, the red dashed line represents the regression line derived from regressing the DNAm age against the chronological age.

Similarly, we also trained a non-cerebellar cortical clock with several non-cerebellar cortical tissues, and the input CpG set was limited to 201 shared age-related CpGs. The resulted model, named CortexClock_common_, performed well for those included training tissues (Supplementary Figure 2). We then applied it to the clean cerebellum dataset (n=752) we collected. As expected, all the cerebellum samples were hugely underestimated by CortexClock_common_. Moreover, the increasing deviations from their chronological age values and the lower than 1 slope value of the regression line (slope=0.54) derived from regressing the predicted DNAm age against the chronological age indicate that the cerebellum ticks at a slower rate than those other brain cortex tissues.

Overall, we arrive at the same conclusion from the two oppositely directed analyses—the cerebellum has a smaller ageing tick rate when measured with the same set of CpGs which were selected as age-associated in both CBL and MTG

### 2.6 How the cerebellum is predicted to age slowly

We then sought to understand the underlying reasons why the cerebellum clock (CerebellumClock_common_) overestimated the non-cerebellar brain tissues and the non-cerebellar brain clock (CortexClock_common_) underestimated the cerebellum tissue. Firstly, comparing the overall methylation levels, the cerebellum has a significantly lower median methylation level than MTG (Figure 5a) and it also has the lowest median methylation level among the five tissue types included in GSE59685 (Figure 5b). When grouping all CpGs into four genomic categories, i.e. island, open sea, shelf and shore, the mean (Figure 5c) and median (Figure 5d) methylation analysis both agreed that the cerebellum is less methylated in the island and the shore. Thus, it is reasonable to conclude that the overall lower methylation level in the cerebellum is due to its lower methylation level in the island and the shore. It should be noted we did not detect any significant correlations between median methylation level change with age in any tissue types or the four genomic categories (Supplementary Figure 3 and 4), indicate the overall lower methylation level in the cerebellum is not due to a different ageing rate.

**FIGURE 5.**
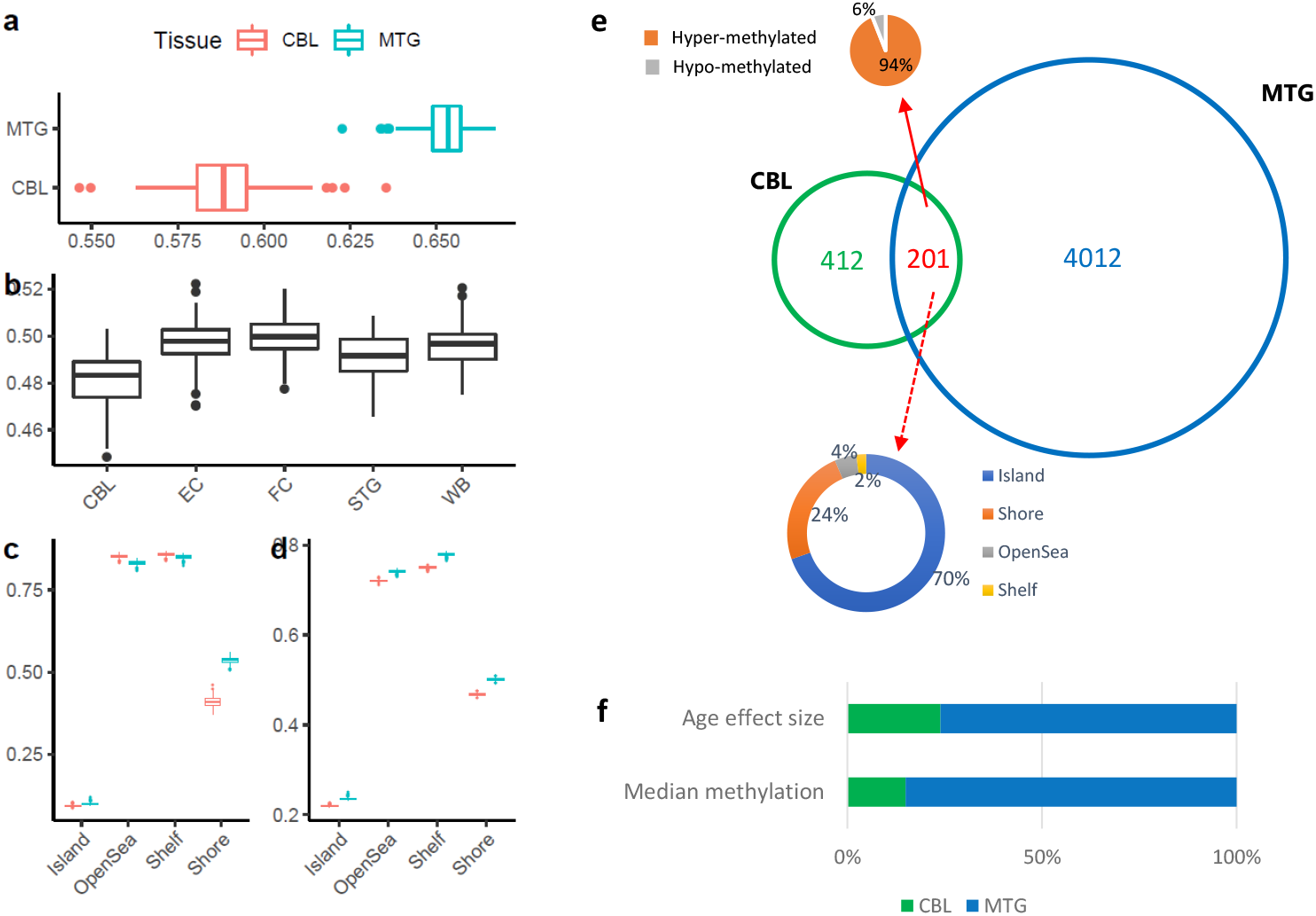
Boxplots illustrate cerebellum has a lower median overall methylation level than (**a**) MTG or (**b**) other four tissues (i.e. EC, FC, STG and WB). (**c**) Median and (**d**) mean methylation comparison both agree the cerebellum has a lower methylation level in the CpG island and shore. Among the 201 common age-associated CpGs, 70% of them are located on the island and 24% of them are located on the shore. The barplot in (**f**) shows the proportions of CpGs, which have a higher age effect sizes or higher median methylation levels in CBL and MTG.

As for the 201 common age-associated CpGs, we have shown that 94% (189) of them gain methylation with age, in other words, their methylation changes positively correlated with age. Interestingly, there are 140 CpGs on the island and 48 CpGs on the shore, and they accounted for 93.5% of the 201 common CpGs (Figure 5e). We have shown that the cerebellum has lower overall mean and median methylation than MTG, as for the common 201 CpGs, the majority of them (85%, 171) also have lower mean/median methylation levels in the cerebellum (Figure 5f). More importantly, more than three-quarters of the 201 CpGs turned out to have a larger age effect size in the MTG when regressing the beta values against age, sex and batches (Figure 5f), meaning those CpGs have higher rates of age-associated methylation change in the MTG than the cerebellum. Collectively, those findings explain why CerebellumClock_common_ not only systematically overestimated the non-cerebellar brain tissues (intercept=11), but also with the overestimation effect increasing with age (slope=1.2). Similarly, they also explain why the cerebellum samples were systematically underestimated by the CortexClock_common_.

## 3 DISCUSSION

In order to test the claim that the cerebellum ages slower, we collected a large set of cerebellum samples (N=756) and assessed their DNAm ages from six representative clocks, including the Horvath’s multi-tissue clock. The results demonstrated that almost all cerebellum samples were severely underestimated by these six representative clocks. This is consistent with previous reports [20, 29]. However, we should not conclude that the cerebellum ages slower only based on these results, as the underestimations may largely reflect the improper usage of DNAm clocks. We found the underestimations were much more severe with the four clocks that were trained with no brain-related tissues. Different tissues may have distinct DNA methylation profiles, the dynamic changes of their methylomes in response to ageing also vary [14]. Horvath’s multi-tissue clock produces relatively accurate age predictions for many vast different tissue/cell types [4], but there is no evidence or guarantee to claim that it has captured the intrinsic mechanism that drives the DNAm changes across the whole body. We do not think it is justified to compare the ageing rates of different tissues by simply comparing their DNAm ages derived from the multi-tissue clock.

By performing age EWAS on the cerebellum, we found a group of (613) significant age-associated CpGs in this tissue. Combined with the penalised linear regression algorithm and a large training dataset (N=752), we built a highly accurate cerebellum specific age clock (CerebellumClock_specific_, r=0.941, MAE=3.18 years). Those results demonstrate the strong and consistent ageing effect in the DNA methylome of the cerebellum. Our age EWAS results show that the cerebellum has a much smaller number of age-associated CpGs; meanwhile there is only a small proportion of age-related CpGs shared between the cerebellum and the MTG, which together indicate a unique pattern of age-related responses in the cerebellar methylome. Among the 201 shared age-associated CpGs, 193 of them gain methylation along with ageing, and the majority of them are less methylated in the cerebellum, which adequately explains why the cerebellum age clock (CerebellumClock_common_) trained on the 201 common CpGs systematically overestimates the MTG samples. As most of the 201 CpGs have lower rates of age-associated changes in the cerebellum compared to MTG, the overestimation effect for the MTG samples increases with age. Our finding agrees that the cerebellar methylome is more resistant to change with ageing, however, we should be cautious about whether this can be translated to a conclusion that cerebellum is biologically younger than other human tissues. To put it simply, if we find one specific CpG site which robustly accumulates methylation from 0% to 100% across a dozen of tissue types in human beings when they grow from birth to 100 years old, then can we conclude that a new tissue type is biologically younger if it only gains methylation from 0% to 50% at that one locus during the same period?

It should be also noted that the above comparisons of DNAm ageing rates between cerebellum and MTG are based on the clocks trained on the 201 shared CpGs. In fact, there are more than twice the number of CpGs found to be age-associated in the cerebellum and even more in MTG. When we apply the cerebellum specific clock (CerebellumClock_specific_), which was trained by using all age-associated CpGs in the cerebellum, in predicting DNAm ages of other brain tissues, we could no longer find a systematic overestimation across all age groups, instead only the individuals aged below 60 years old were overestimated, by contrast, the above 60 years old group was clearly underestimated. We conclude that this is due to the improper usage of the clock, as the CerebellumClock_specific_ consists of many cerebellum-specific CpGs.

It is easy to understand the DNAm age comparison between samples from the same tissues, i.e. we are confident the sample A is biologically younger than sample B, when DNAm age of sample A is much smaller than sample B and they are from the same tissue. However, we still lack sufficient evidence to confidently compare the biological ages between samples from different tissues. For example, as recently reported by Jonkman and colleagues, the Horvath’s multi-tissue clock predicts naive T cells to be up to 30 years younger than activated T cells from the same donor [41]. Can we conclude the naive T cells are biologically 30 years younger than activated T cells? Similarly, when predicted by our CerebellumClock_specific_, the non-cerebellar brain tissues are predicted to be at least 11 years older than the cerebellum (Figure 4a), however we can not claim that those non-cerebellar brain tissues are biologically 11 years older than the cerebellum, as we could easily find one CpG or several CpGs combined that distinguishes the cerebellum from other brain tissue, then add it to the existing model and assigns it with a coefficient to counteract the 11 years gap. Then the new adjusted clock should not produce DNAm age predictions with systematic huge differences between the cerebellum and other brain tissues. As proposed by Liu et al., the many non-age-related CpGs in Horvath’s multi-tissue clock [4] may actually be reflecting and adjusting for tissue differences [39].

Another angle for ageing rate comparisons is to look at the Telomere Length (TL) shortening rates. Telomeres are protective DNA-protein complexes at the termini of chromosomes [42] and telomere attrition is considered an important hallmark of human ageing [1]. As comprehensively studied by Demanelis et al. [43], the average relative TL (RTL) varies across different tissue types, for instance, the average RTL is the lowest in the whole blood and the longest in testis. Even though they found TL can shorten at different rates with ageing between several tissue types, the majority tissues do not show significant difference in age-dependent shortening rates, and there is no evidence to claim that different tissue types age at rates proportional to their TL shortening rates.

We should acknowledge some limitations of this study. First, due to the scarcity of cerebellum samples, the majority of our collected cerebellum samples are from elderly individuals aged above 60 years old. It would be very valuable to test our hypothesis that the Horvath’s multi-tissue clock would overestimate most of the cerebellum samples from young individuals aged below 30 years old. Second, our age EWASs on the cerebellum and MTG were also based on a very elderly population which has a relatively narrow age range, as demonstrated by Vershinina and colleagues [44], lots of age-associated CpGs do exhibit nonlinear methylation changes with age. Thus our age EWASs may have missed many CpGs that are strongly age-associated in the younger age group but to be a much-attenuted association in the aged group. Future studies that include more young individuals should reveal a more complete picture of age-associated changes of the cerebellar methylome.

## 4 METHODS

### 4.1 DNAm datasets

The DNAm samples were collected from the public data repository—Gene Expression Omnibus (GEO). The cerebellum samples are from six datasets, including GSE134379 [32], GSE59685 [34], GSE105109 [36], GSE125895 [35], GSE61431 [37] and GSE72778 [38]. They were included according to the following criteria: contains at least 20 cerebellum samples; with age annotations; and raw IDAT files or methylated and unmethylated intensity files are available. The cerebellum samples were used to reveal the underestimation issues for the cerebellum tissue by six representative clocks and were also used to train cerebellum age clocks. Apart from cerebellum samples, GSE134379 [32] also includes DNAm microarray data of middle temporal gyrus from the same 404 individuals, thus it was used to perform age EWASs on the two brain tissues. GSE59685 [34] includes 531 DNAm samples of five tissues, i.e. cerebellum, entorhinal cortex, frontal cortex, superior temporal gyrus and whole blood, from donors (N = 122) archived in the MRC London Brainbank for Neurodegenerative Disease. GSE59685 and GSE134379 were also used to compare the DNAm ages of different tissues which were estimated by our trained cerebellum clocks. The DNAm samples of the non-cerebellar brain tissues in four datasets, i.e. GSE134379 [32], GSE74193 [45], GSE80970 [46] and GSE61431 [37], were used to train the cortex age clocks, the four datasets were selected to ensure a relatively uniform sample distribution across all age groups in the adult population.

### 4.2 Data preprocessing

For all the DNAm datasets, after downloading from the GEO, they were read into R by using the *iadd2* function from the ‘bigmelon’ package [47] when raw IDAT files were available. For those datasets in which only text formatted intensity files exist, the methylated and unmethylated intensities were extracted and read into R directly. Then the raw methylation beta values is calculated as: 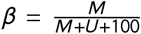, where *M* denotes methylated intensities and *U* denotes unmethylated intensities. For all those samples, we estimated their sex by using the *estimateSex* function [48] from the wateRmelon package [49], any samples with mismatches between its reported sex and the estimated sex from the DNAm data were excluded for downstream analysis. Also, the beta value density distributions of samples within each dataset were manually checked to remove any samples with abnormal distribution profiles.

### 4.3 DNA methylation age prediction

The DNAm age prediction of the six representative clocks, i.e. Hannum2013 [3], Horvath2013 [4], Horvath2018 [27], Levine2018 [23], Zhang2019 [22] and Shireby2020 [16], was completed by using the *methyAge* function from the ‘dnaMethyAge’ R package [50]. Only methylation beta values are required to feed into the *methyAge* function. Note, when calculating the DNAm age of Horvath2013, the raw beta values are firstly normalised with an adjusted BMIQ which has a fixed reference, this is consistent with Horvath’s original publication [4]. To calculate the DNAm age of Zhang2019, the beta values of each sample are first subjected to Z-score normalisation [22]. For the remaining clocks, no normalisation steps were applied. The difference between DNAm age and chronological age are measured as:

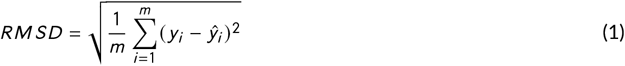

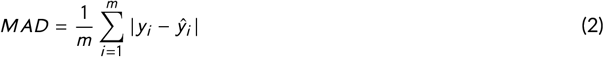

where *y_i_*, represents the chronological age of the *i_th_* sample, 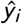 represents the predicted DNAm age of the *i_th_* sample, *m* denotes the number of all samples. RMSD: root mean squared deviation; MAD: mean absolute deviation.

### 4.4 Epigenome-wide association study

The age EWASs were performed on GSE134379 [32] which includes DNAm microarray data of two brain tissues (CBL and MTG) in every individual from a large elderly population (N=404). The CBL samples and MTG samples were normalised by the *adjustedDasen* [51] from the ‘wateRmelon’ package [49] separately. These probes target CpGs mapped to sex chromosomes or reported to have cross hybridizing issues were removed from downstream analysis [52]. To find out age-associated differentially methylated CpGs across the genome in the two brain tissues, we fitted the following linear regression model for each CpG site involved in the two tissues separately:

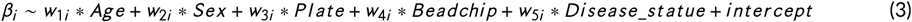

where *β_i_*, is the methylation beta value of the *i_th_* CpG, *w*_1*i*_, is the coefficient of chronological age for the *i_th_* CpG. The *t statistic* of the coefficient *w*_1*i*_ is checked in the *Student’s t* distribution to determine the p-value. After that, the p-values of all studied CpGs were adjusted with the Benjamini & Hochberg method. A CpG is called to be significant age-associated when its adjusted p-value (or FDR) less than 0.01.

### 4.5 The construction of DNAm clocks

Prior to any training steps, all DNAm samples were normalised by a modified version of *adjustedDasen* [51] method from the ‘wateRmelon’ package [49], in which the modified *adjustedDasen* is supplied with a fixed reference to reduce the batch variance between different datasets. Also, the chronological age is log-transformed.

The new clocks mentioned in this study were all trained by the penalised linear regression algorithm—Elastic net [40], which is essentially a linear combination of the L1 and L2 penalties of the lasso regression and ridge regression.

The loss function of Elastic net is defined as:

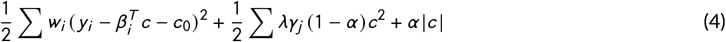

where the *β_i_*, denotes the methylation beta value of *ith* CpG, *c* is the coefficient vector of all the CpG accounted, *α* is the critical parameter that controls the weights of the L1 and L2 penalties and has been defined prior to the training.

We used the *cv,glmnet* function from the ‘glmnet’ R package [53] to train the Elastic net models. To train the CerebellumClock_specific_, the input samples are the 752 cerebellum samples from six independent datasets, the input CpG set of each sample was restricted to the 613 age-associated CpGs in the cerebellum, alpha was set to 0.5, and 10fold cross-validation was used to determine the optimal coefficient combination. We made use of leave-one(dataset)-out cross-validation to infer the age prediction performance of the CerebellumClock_specific_. Specifically, we have six independent cerebellum datasets, then for each round of the total six rounds of cross-validation process, one dataset was taken out and their DNAm ages were estimated by the model trained on the remaining five datasets, after six rounds, the DNAm ages of samples from the six datasets were derived and they were not overfitted by the training process. In the same way, the CerebellumClock_common_ was trained on the same 752 cerebellum samples but the input CpG sets were restricted to the 201 shared age-associated CpGs. Another difference was the alpha value was set to 0.2 to let the final model includes more CpGs from the 201 CpGs.

The training of CortexClock_common_ also employed Elastic net linear regression, the training samples were those of non-cerebellar brain tissues from four independent datasets, the input CpG set of each sample was also restricted to the 201 shared age-associated CpGs and the alpha was set to 0.2. As we only have four separate datasets, and only GSE74193 [45] has a wide age range, we employed a 10-fold cross validation to measure the age prediction performance of CortexClock_common_. That is to say, we first randomly separate all the training samples into equal 10 portions, for each round of the total 10 rounds of cross-validation processes, we took one portion out and their DNAm ages were then estimated by the model trained on the remaining 9 portions. After 10 rounds, the DNAm ages of samples from all ten portions are obtained and they were not overfitted by the training process.

The coefficients of involved CpGs in each model are listed in Supplementary 1.

### 4.6 Software

All the analysis were conducted in R (version 3.6.0) [54] under Linux environment. The scatter plots in Figure 1,3,4 were produced by the *getAccel* function with proper settings from the ‘dnaMethyAge’ R package [50]. The three constructed models of CerebellumClock_specific_, CerebellumClock_common_ and CortexClockcommon are readily available to be applied in independent DNAm samples by calling the *methyAge* function from the ‘dnaMethyAge’ R package [50] with the ‘clock’ parameter setting as ‘Cerebellum_specific’, ‘Cerebellum_common’ and ‘Cortex_common’ respectively. GO analyses were conducted using the *gometh* function in the ‘missMethyl’ package [55] which tests gene ontology enrichment for significant CpGs while accounting for the differing number of probes per gene present on the 450k.

## Funding and Acknowledgements

YW was supported by a University of Essex Doctoral studentship. LCS was supported by MRC grants [MR/W004984/1, MR/R005176/1]. KDM is funded by the Economic and Social Research Council [ES/M010236/1] and the Engineering and Physical Sciences Research Council [EP/P017487/1, EP/R02572X/1, EP/V000462/1]. XZ is funded by the Engineering and Physical Sciences Research Council [EP/V034111/1]. The authors acknowledge the use of the High Performance Computing Facility (Ceres) and its associated support services at the University of Essex in the completion of this work.

## Author Contributions

YW conceived the study. YW collected the data, performed most analyses except for the GO analyses. OAG performed the GO analyses and summarised this part. LCS and XZ contributed to the interpretation of the results. YW drafted the manuscript with critical contributions from LCS, KDS, OAG and XZ. XZ, KDM and LCS advised and oversaw the work. All authors read and approved the final manuscript.

## Supporting Information

Additional supporting information may be found online in the Supporting Information section at the end of the article.

